# Dynamical consequences of regional heterogeneity in the brain’s transcriptional landscape

**DOI:** 10.1101/2020.10.28.359943

**Authors:** Gustavo Deco, Kevin Aquino, Aurina Arnatkevičiūtė, Stuart Oldham, Kristina Sabaroedin, Nigel C. Rogasch, Morten L. Kringelbach, Alex Fornito

## Abstract

Brain regions vary in their molecular and cellular composition, but how this heterogeneity shapes neuronal dynamics is unclear. Here, we investigate the dynamical consequences of regional heterogeneity using a biophysical model of whole-brain functional magnetic resonance imaging (MRI) dynamics in humans. We show that models in which transcriptional variations in excitatory and inhibitory receptor (E:I) gene expression constrain regional heterogeneity more accurately reproduce the spatiotemporal structure of empirical functional connectivity estimates than do models constrained by global gene expression profiles and MRI-derived estimates of myeloarchitecture. We further show that regional heterogeneity is essential for yielding both ignition-like dynamics, which are thought to support conscious processing, and a wide variance of regional activity timescales, which supports a broad dynamical range. We thus identify a key role for E:I heterogeneity in generating complex neuronal dynamics and demonstrate the viability of using transcriptional data to constrain models of large-scale brain function.

---

Brain dynamics are often described as complex, displaying properties that are interposed between order and disorder (*1, 2*). These complex dynamics arise from two cardinal factors: (1) the properties of local population activity within each brain region; and (2) the mutual influences that these populations exert on each other, as mediated by their interconnecting anatomical projections.

Biophysically plausible non-linear models of brain dynamics have been used to clearly demonstrate the importance of anatomical connectivity in supporting complex patterns of network activity. For example, when simulated on human inter-regional structural connectivity (SC) matrices (connectomes) generated with diffusion magnetic resonance imaging (MRI), these models can reproduce empirically observed patterns of inter-regional functional connectivity (FC) (reviewed in (*2*)). In macaque, a biophysical model simulated on a connectome derived from viral tact tracing indicated that both feedback projections and a heterogeneous distribution of connection strengths across projections are required for the emergence of ignition-like dynamics (*3*), which are thought to be a necessary precondition for conscious processing (*4*).

By comparison, variations of local circuitry within each region have received less attention. Precise coordination of firing across spatially distributed neuronal ensembles depends on a delicate balance between excitatory and inhibitory activity. The first wave of biophysical models of large-scale brain activity treated all local population dynamics as homogenous, driven by sub-populations of inhibitory and excitatory neurons with uniform properties across all brain regions (*5*–*7*). However, heterogeneity in regional cytoarchitecture has been noted for over a century, as famously mapped by Brodmann (*8*), and subsequent work has provided mounting evidence for large-scale cortical gradients in gene expression, cellular composition, connectivity, and function (*9, 10*). Thus, quantitative variations of local circuit properties across the cortex also play an important role in shaping complex neural dynamics, likely through their influence on the local ratio, or balance, of excitatory and inhibitory cell activity (E:I balance) (*9*).

Ideally, one could model regional heterogeneity using empirically observed estimates of areal variations in excitatory and inhibitory cell counts, or a related measure of cell-type specific activity. No such data are available on a large scale for primate brains, but some early modelling work has used alternative methods to constrain biophysical models. Deco et al. (*11*) showed that heterogeneity in local feedback inhibition, in which E:I balances were algorithmically adjusted to achieve a uniform firing rate of 3 Hz across all regions (in line with experimental data; (*12*–*14*), yields networks with more stable dynamics and improves model fits to static measures of FC when compared with a strictly homogeneous model. Using data in macaque, Chaudhuri et al. (*15*) scaled the excitatory input strength of each of 29 cortical areas according to its assumed hierarchical position, as determined by laminar patterning of inter-regional projections (*16*). They found that this heterogeneity was essential for generating dynamics characterized by a hierarchy of regional activity timescales, in which areas with a higher hierarchical position show a prolonged response to a simulated sensory input. This result mirrors previous single unit findings in macaque cortex (*17*) and aligns with other evidence for regional differences in activity timescales (see also (*18*–*20*)). Two subsequent studies modelling human functional MRI found that, when compared to a regionally homogeneous model, scaling a region’s excitatory activity according to its assumed hierarchical position improves fits to measures of edge-level and node-level functional connectivity, more faithfully reproduces empirically observed regional variations in activity timescales, and better explains FC disturbances in people with schizophrenia (*21, 22*). A complementary study used a stochastic optimization procedure to invert local dynamical parameters such as recurrent synaptic connectivity strengths to optimally fit functional MRI data (*23*). Regional variations in the estimated local dynamical parameters showed hierarchical ordering and correlated with variations in myelin and layer-specific neuronal density, presumed cognitive functions (derived from meta-analysis of task-based fMRI studies), and a principal gradient of resting-state FC (*24*).

These initial models of regional heterogeneity demonstrate the critical role of local variations in E:I balance for generating realistic patterns of brain-wide dynamics. However, in the absence of a gold-standard biological constraint, there has been considerable variability in how heterogeneity has been instantiated within a given model. There is also substantial variability in the dynamical properties that have been modelled. For example, modelling work in macaques shows that it is, in principle, possible to find a parameter regime that results in ignition-driven and hierarchical dynamics (*3, 15*), but the parameters were not fitted to empirical data making it difficult to evaluate the model’s capacity to reproduce other key features, such as realistic patterns of static or dynamic FC. Similarly, studies in humans have modelled specific outcome measures, such as static or dynamic FC at the level of individual connections (*6, 7, 22, 25*), the mean static FC of each region (*22*), or timescale hierarchies (*3, 22*), meaning that it is unclear whether the working point for a model fitted to one property can also capture other properties. Developing a coherent approach that can account for each of these diverse features is an essential step towards developing a unified model that can parsimoniously explain diverse empirical properties of complex whole-brain dynamics.

Here, we seek to address this aim by introducing a model that leverages brain-wide transcriptional data (*26*) to constrain regional heterogeneity and tune each region’s dynamics according to regionally-specific measures of inhibitory and excitatory receptor gene expression. We evaluate the performance of this model with respect to some of the major outcome measures evaluated thus far in the literature; namely, edge-level static FC, node-level static FC, FC dynamics (FCD), ignition capacity, and timescale hierarchy. We show that our model more faithfully reproduces empirical properties of static and dynamic FC than other models in which heterogeneity is imposed using plausible biological constraints, such as regional variations in the ratio of T1-to-T2-weighted MRI signal (T1w:T2w) and loadings on the dominant mode of gene expression (*22, 27*). We then show that transcriptomically-constrained models show the greatest capacity for ignition-like dynamics and display a broad range of regional activity timescales. Our results indicate that regional variations in the transcriptional activity of inhibitory and excitatory receptor genes provide a viable means for constraining biophysical models of large-scale neural dynamics and, when coupled with an empirically-derived connectivity matrix, can offer a parsimonious account of diverse properties of large-scale human brain activity.

## Results

### Overview

Our overall analysis strategy is outlined schematically in Figure 1 and the details of our approach are provided in the Methods. Briefly, we use diffusion MRI to generate a group-level representative structural connectome for a sample of 293 healthy individuals, representing SC between each pair of 68 regions defined according to an extensively used neuroanatomical atlas (Figure 1A). We also estimate, for a partially overlapping sample of 389 individuals, various empirical FC properties using task-free resting-state functional MRI to probe spontaneous blood-oxygenation-level-dependent (BOLD) dynamics (Figure 1B).

**Figure 1.**
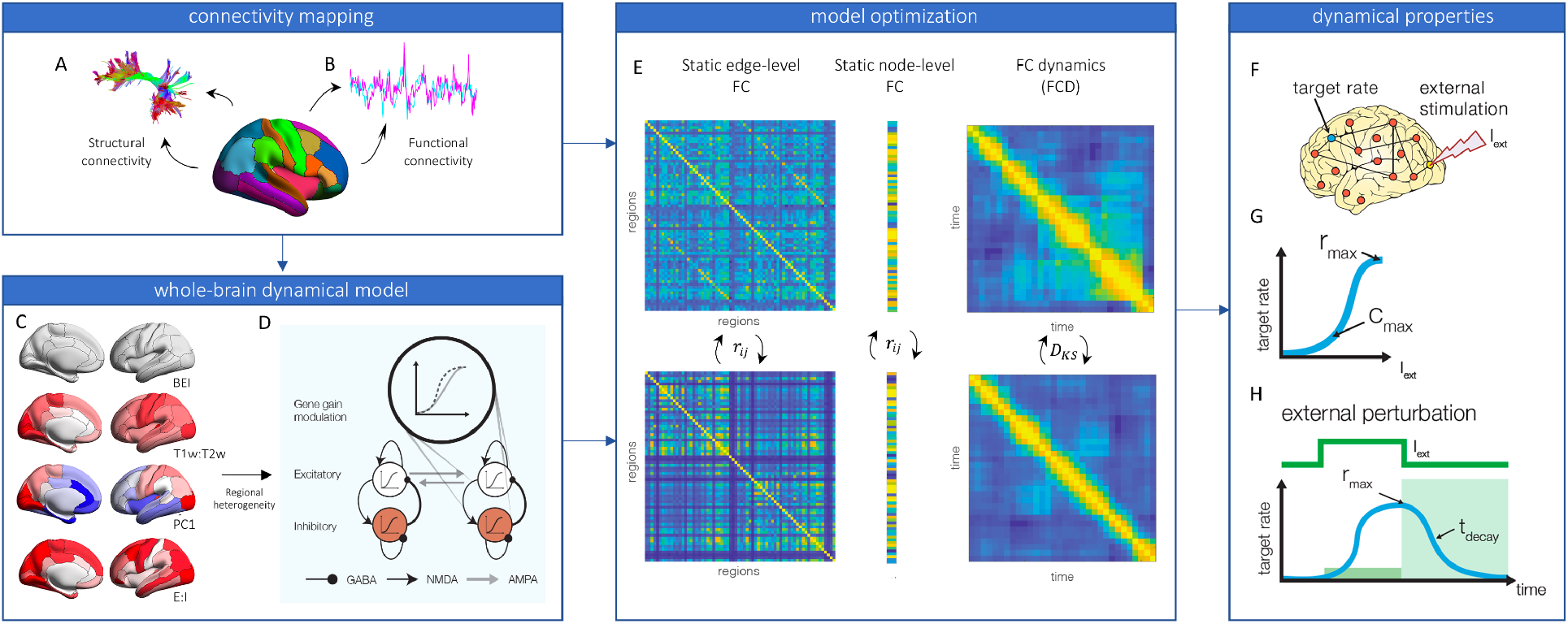
Workflow for model fitting and evaluation. We use empirical diffusion and functional MRI data to generate sample-averaged structural connectivity (**A**) and functional connectivity (**B**) matrices, respectively. The structural connectivity matrix, and different forms of regional heterogeneity (**C**), are used to respectively define the coupling between, and modulate the gain of, neuronal populations within a large-scale balanced excitation-inhibition (BEI) dynamical model (**D**). Panel C shows the different forms of regional heterogeneity used. The standard implementation of the BEI model assumes uniform gain across all regions (top). Empirical estimates of T1w:T2w, the primary mode (PC1) of gene expression, and the ratio of excitatory to inhibitory receptor expression (E:I), are used to modulate the population gain function in the heterogenous models. (**E**) For each model, the global coupling parameter, *G*, is tuned to optimize the fit to three empirical FC properties: static edge-level FC (left), static, node-averaged FC (middle), and FC dynamics (FC; right). Gain parameters *B* and *Z* are then tuned to further improve fits to these properties. **F:** After finding the optimal working point for each model, we simulate a focal perturbation of activity in occipital cortex and measure two dynamical properties of the network: ignition and decay timescale. **G**: To measure ignition, we characterize the response of each area to focal stimulation as the product of two quantities: the maximum firing rate achieved at the highest stimulation intensity, *r*_*max*_, and the concavity of the regional response function (i.e., the maximal second derivative), *c*_*max*_. The ignition capacity of the model is taken as the average of these regional values. **H:** To measure the intrinsic timescale of each region, we fit an exponential curve to each region’s firing rate as it returns to baseline after termination of the maximal stimulation intensity (**H**).

We incorporate different forms of regional heterogeneity (Figure 1C) into a whole-brain model in which the local neuronal dynamics of each brain region evolve according to a dynamic mean field reduction analytically derived from the collective behaviour of empirically validated integrate-and-fire (80% excitatory and 20% inhibitory) neurons (*13, 14*). Regional dynamics in the model are driven by a pool of excitatory neurons, a pool of inhibitory neurons, the interactions between them, and with the net output of other areas, as mediated by the SC matrix (Figure 1D). In this model, local feedback inhibition is adjusted separately for each region to ensure a consistent firing rate of 3 Hz across regions (*11*). We thus refer to this model as the balanced excitation-inhibition (BEI) model. Although the local adjustments in this model introduce some degree of regional heterogeneity, the firing rates are constrained to be uniform across regions so we consider this BEI model as a homogeneous benchmark against which we evaluate more sophisticated heterogeneous models that allow intrinsic dynamical properties to vary across regions (see (*22*) for a similar approach). More specifically, the activity levels of excitatory and inhibitory populations in each region, *i*, of the BEI model are given by independent sigmoid functions, regulated by a single gain parameter *M*_*i*_. In our benchmark BEI model, this parameter is set to a fixed value for all regions, imposing uniform excitability across brain regions (Figure 1C).

We compare the performance of this homogeneous BEI model to three different heterogeneous models. In the first, we use an approach similar to (*22*) and adjust *M*_*i*_ according to variations in the regional mean T1w:T2w, which has been proposed as a proxy for cortical hierarchy (*27*). In the second heterogeneous model, we adjust *M*_*i*_ according to the first principal component (PC1) of transcriptional activity for 1,926 brain-specific genes (*27*) that survived our quality control criteria, as quantified in the Allen Human Brain Atlas (AHBA) (*26, 28, 29*). The first PC of these genes correlates with the T1w:T2w ratio and other measures of cortical hierarchy (*27*). In the final model, we use a more hypothesis-driven approach and adjust *M*_*i*_ according to variations in the ratio of regional AHBA expression values for genes specifically coding for AMPA, NMDA, and GABA receptors (see Methods). We refer to this model as the E:I model. Cortical surface renderings displaying regional variations in T1w:T2, PC1, and E:I values are shown in Figure 1C.

In all three heterogeneous models, we assume a linear scaling between the regional biological measures of heterogeneity, *R*_*i*_, and the effective gain within a region (*22*), given by *M*_*i*_ = 1 + *B* + *ZR*_*i*_, where the two unknown free parameters correspond to a bias, *B*, and a scaling factor, *Z* (see *Methods*). Studying how these two free parameters affect the global dynamics of the model allows us to investigate the role of regional heterogeneity.

For the homogeneous BEI and three (T1w:T2w, PC1, E:I) heterogeneous models, we assume that all diffusion MRI-reconstructed streamline fibres have the same conductivity and thus the coupling between different brain areas is scaled by a single global parameter, *G*. We first tune the *G* parameter of the BEI model to adjust the strength of effective coupling in the model and identify the brain’s dynamic working-point by fitting the model to three empirical FC properties (Figure 1E): (1) the Pearson correlation between model and empirical estimates of static (i.e., time-averaged) FC estimated across all pairs of brain regions (more specifically, the correlation between the values in the upper triangles of the model and empirical FC matrices); (2) the Pearson correlation between model and empirical node-level estimates of average FC (*22*); and (3) similarity in functional connectivity dynamics (FCD), estimated as the distance between model and empirical distributions of frame-by-frame FC properties (as quantified using the Kolmogorov-Smirnov distance, *D*_*KS*_)(*25*).

After finding the working point of each model, we evaluate its dynamical properties in two ways. First, we consider the ignition capacity of the model, quantified by examining how activity propagates through the network following a simulated focal perturbation (Figure F-G). Second, we examine regional heterogeneity in intrinsic timescales, quantified by the signal decay rate of regional activity following a simulated focal perturbation (Figure F,H). Our approach thus allows us to determine the performance of each model in capturing diverse dynamical phenomena at a single working point.

### Fitting the Homogeneous Model

We first evaluate the performance of the homogeneous BEI model in reproducing empirical properties of resting-state FC data. We determine and fix the global coupling parameter, *G*, with all regional gain parameters, *M*_*i*_ = 1 (i.e. heterogeneity is not considered). As done previously (*11*)(*30*), we examine how well the model fits, as a function of *G*, three different properties of empirical resting-state fMRI recordings: edge-level static FC, FCD, and node-level FC (see Figure 1B and Methods for further details). The results of this analysis are shown in Figure 2A-C. In all cases, we consider group-averaged values for both empirical data (across subjects) and the model (across an equivalent number of simulated trials with the same duration as the human experiments) (see *Methods*). Our results align with previous investigations to clearly show [40, 44] that fitting FCD, which captures the rich spatiotemporal structure of the fMRI data, is a stronger constraint on the model. Indeed, where static edge and node FC fits are consistently high across a broad range of *G*, FCD yields a clear global optimum at *G* = 2.1.

**Figure 2.**
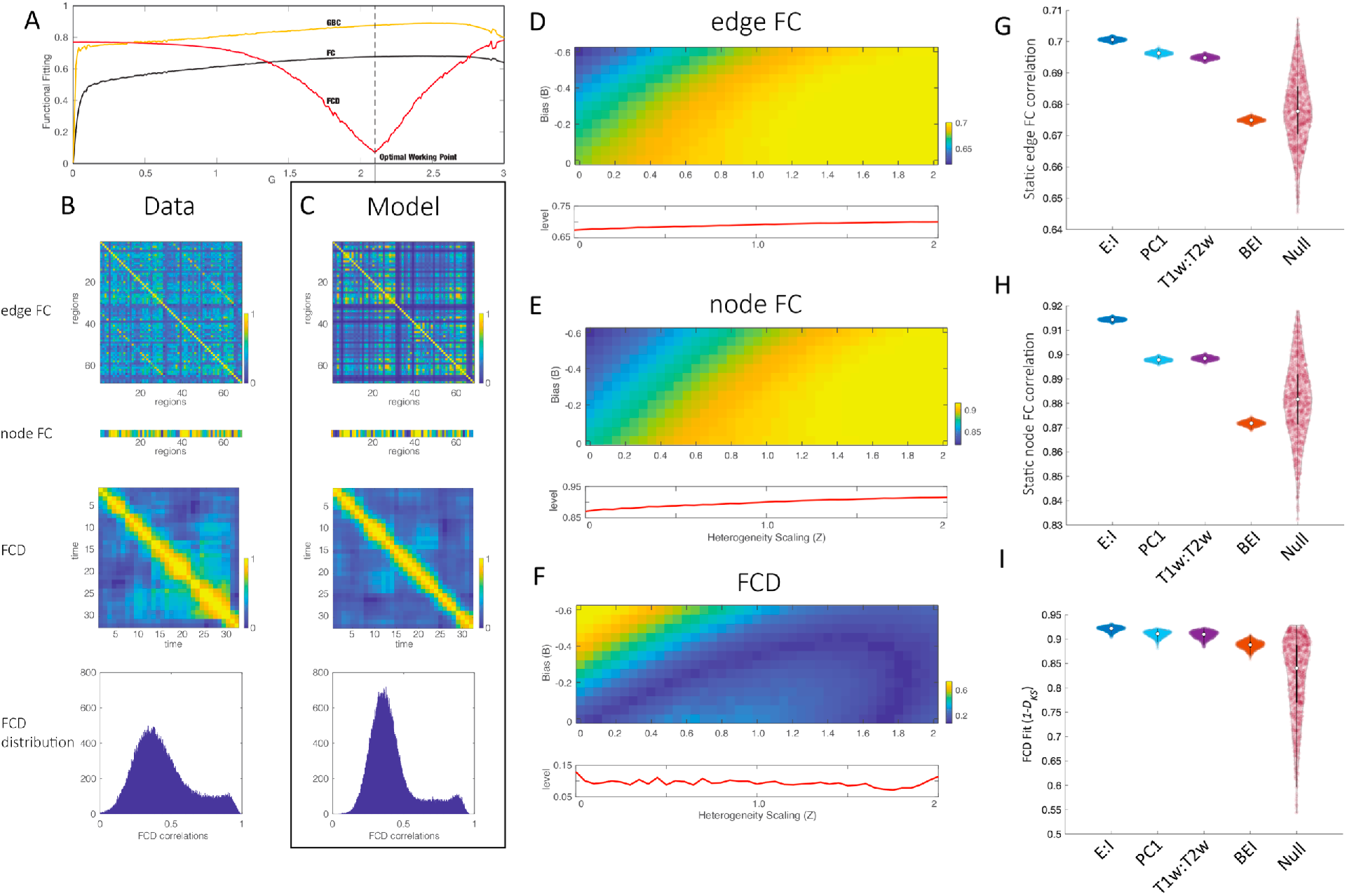
Model optimization and evaluation. The homogeneous BEI model is first fitted to the empirical data. **A:** The correlation between model and empirical static edge-level and node-level FC, as well as the *D*_*KS*_ between empirical and model FCD distributions, as a function of the global coupling parameter, *G*. Only the fit to FCD shows a clear optimum at *G* = 2.1 (note that lower FCD fit values indicate better fit). Panels **B** and **C** show the FC matrix, node-level average FC values, FCD matrices, and FCD distributions for the empirical data (**B**) and model (**C**). Using *G* = 2.1, we introduce regional heterogeneity (Figure 1C) and further optimize the two gain parameters *B* and *Z*. Panels **D, E**, and **F** show the parameter landscapes for edge-level FC, node-level FC, and FCD, respectively, for the E:I model. Below each 2D space we show the 1D graph of the level heterogeneity scaling (*Z*) for the optimal bias parameter found as the one corresponding to the optimum FCD fitting (see Methods). This procedure is repeated for each heterogeneous model to find its optimal working point. Lower values indicate better fit in panel **F**. Panels **G, H, I** compare the performance of the homogeneous BEI model and heterogeneous T1w:T2w, PC1, E:I, and null expression models according to the edge-level FC, node-level FC, and FCD benchmarks, respectively. Distributions are for values obtained over multiple runs of the optimum models (see Methods). We harmonized the fit statistics so that higher numbers indicate better fit (i.e., for the case of FCD, where a low *D*_*KS*_ indicates better fit, we show 1 − *D*_*KS*_). The E:I model shows the best performance in all cases.

### Introducing regional heterogeneity

We now study how regional heterogeneity affects the fitting of static FC, GBC, and FCD. Spatial maps for each form of biological heterogeneity used in our modelling are shown in Figure 1C. The T1w:T2w and PC1 maps are strongly correlated (*ρ*= 0.82, *p*_*spatial*_ < .001), as shown previously (*27*), whereas the PC1 and T1w:T2w maps show weaker correlations with the E:I map (*ρ* = 0.40, *p*_*spatial*_ = .005 and *ρ* = 0.39, *p*_*spatial*_ < .011, respectively). This result indicates that each spatial map introduces a different form of biological heterogeneity to the benchmark BEI model. We introduce this heterogeneity by modulating the regional gain functions *M*_*i*_, at the optimal working point of the homogeneous BEI model (*G* = 2.1), through the bias and scaling parameters introduced above, denoted, *B* and *Z*, respectively. Critically, we exhaustively search a broad range of values for *B* and *Z* to comprehensively map the parameter landscape and select the corresponding optimal working point for the E:I model (*B* = −0.3; *Z* = 1.8), PC1 model (*B* = −0.75; *Z* = 1.6), and T1w:T2 model (*B* = −0.7; *Z* = 1.4), as shown in Figure 2D-F for the E:I model. This figure shows that, despite some degeneracy in the parameter landscapes, there is a clear regime in which all three empirical properties are fitted well by the model, particularly for *Z* ≈ 1.7.

For these optimal values, we simulate each dynamical model 1000 times to account for the inherent stochasticity of the models and compute the respective measures of model fit. Figures 2G-I show the distributions of fit statistics across runs for the homogeneous and three heterogenous models. In addition, we show results for a null ensemble of models in which the regional E:I receptor gene expression values were spatially rotated (*31, 32*) to generate surrogates with the same spatial autocorrelation as the empirical data. Across all three benchmark properties to which the data were fitted––static FC, GBC, and FCD––the heterogeneous models perform better than the homogeneous model (all *p*_*bonf*_ < .05). We also find a consistent gradient of performance across all benchmarks, with the E:I model performing best, followed by the PC1 and T1w:T2w models, and the homogeneous model showing the poorest performance. For each benchmark metric, the performance of the E:I model was significantly better than all other models (all *p*_*bonf*_ < .05).

### Ignition capacity

We now evaluate additional dynamical properties of each model that cannot be directly evaluated with respect to empirical fMRI data but which are thought to be essential features of complex neural dynamics. We first consider ignition capacity. Ignition refers to the capacity of a sensory input to trigger self-supporting, reverberating, temporary, metastable, and distributed network activity, and is thought to be a necessary condition for conscious perception of the stimulus (*4, 33*). This process is non-linearly related to stimulus intensity, such that evoked activity is confined to local sensory areas for low levels of stimulation (which supports subliminal perception) and then leads to explosive activation of distributed cortical systems once an appropriate threshold is reached.

Figure 1F-G shows the strategy that we follow to quantify ignition capacity. For a given specific working point of the whole-brain model, we compute how the brain broadcasts information after artificially stimulating regions of occipital cortex (namely, the cuneus, lateral occipital, lingual gyrus, and pericalcarine regions in the Desikan-Killiany atlas (*34*)). We measure the evoked responses at the level of population firing rates rather than simulated BOLD signal changes to have direct access to the millisecond timescale. To quantify the effect of occipital stimulation on activity in each of the other 67 brain regions, we plot, for each non-stimulated region, how its population firing rate changes as a function of occipital stimulation intensity. Two quantities are relevant here: (1) the maximum firing rate achieved at the highest stimulation intensity, *r*_*max*_; and (2) the speed with which the firing rate increases beyond a given intensity threshold, *c*_*max*_, which we quantify as the concavity of the regional response function (i.e., the maximal second derivative). We then summarize the ignition capacity of each non-stimulated region *i* as *I*(*i*) = *c*_*max*_(*i*)×*r*_*max*_(*i*), and estimate the global ignition capacity of the brain as the mean across all regions,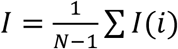.

Figure 3A shows cortical surface renderings of regional *I*(*i*) values obtained following occipital stimulation over 30 runs of the homogeneous BEI and heterogenous E:I models. The occipital stimulation in the E:I model triggers stronger and more widespread ignition than in the BEI model. This observation is confirmed quantitatively in Figure 3B. The E:I model shows higher ignition capacity than all other models (all *p*_*bonf*_ < .05), suggesting that this model is able to rapidly propagate focal perturbations with higher fidelity than in the other models. The PC1 and T1w:T2w models also show significantly greater ignition capacity than both the BEI model and spatial null model, with the BEI model showing significantly lower ignition capacity than the spatial null (all *p*_*bonf*_ < .05). These results indicate that some form of regional heterogeneity is important for ignition-driven dynamics and that the most potent ignition is observed when heterogeneity is constrained by biological measures, particularly regional variations in the transcriptional activity of E:I receptor genes.

**Figure 3.**
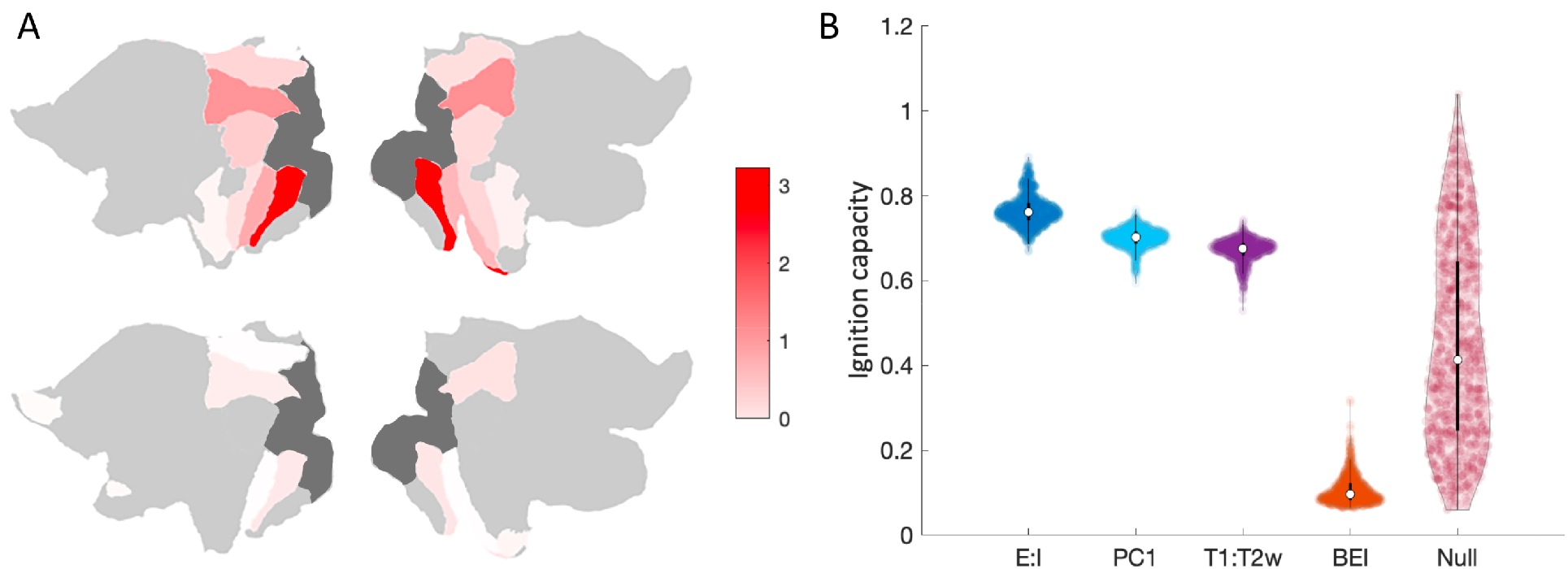
Evaluating the ignition capacity of each dynamical model. **A:** Cortical flat maps of average regional ignition values following stimulation of four regions of occipital cortex (dark grey), observed for the E:I model (left) and BEI model (right). The E:I model shows strong ignition of distributed areas whereas the BEI model shows negligible ignition of non-stimulated regions. **B:** Distributions of global ignition values obtained for different runs of the optimal E:I, PC1, T1w:T2w, BEI, and spatial null models.

### Decay timescales

To characterize temporal hierarchies in the model, we use the same virtual stimulation approach as deployed in our analysis of ignition capacity. Specifically, we virtually stimulate bilateral pericalcarine cortex and quantify the temporal decay of activity in each non-stimulated target area by fitting an exponential curve to its firing rate as it returns to spontaneous conditions after termination of the maximal stimulation intensity (Figure 1H). The value of the exponent can be used to index the intrinsic activity time scale of each brain region and can thus be used to evaluate temporal hierarchies in the brain (*15*) (see Methods).

We first confirm that this measure yields a sensible spatial patterning of regional timescale variations. Figure 4A shows the stimulation responses for the stimulated area and eight non-stimulated regions, approximately ordered according to assumed position in the cortical hierarchy. A clear trend is evident, in which responses are prolonged for higher-order association areas compared to sensory regions, as shown previously (*15, 20, 35*). Figure 4B shows that the activity decay time courses following the termination of stimulation are faster for areas that assume a lower position in the hierarchy. Accordingly, we find a significant positive correlation between regional decay values and an independent measure of cortical hierarchy, T1w:T2w (*ρ*= 0.39, *p*_*spatial*_ = 0.33; Figure 4C), such that higher T1w:T2w is associated with more rapid activity timescales (higher decay exponents).

**Figure 4.**
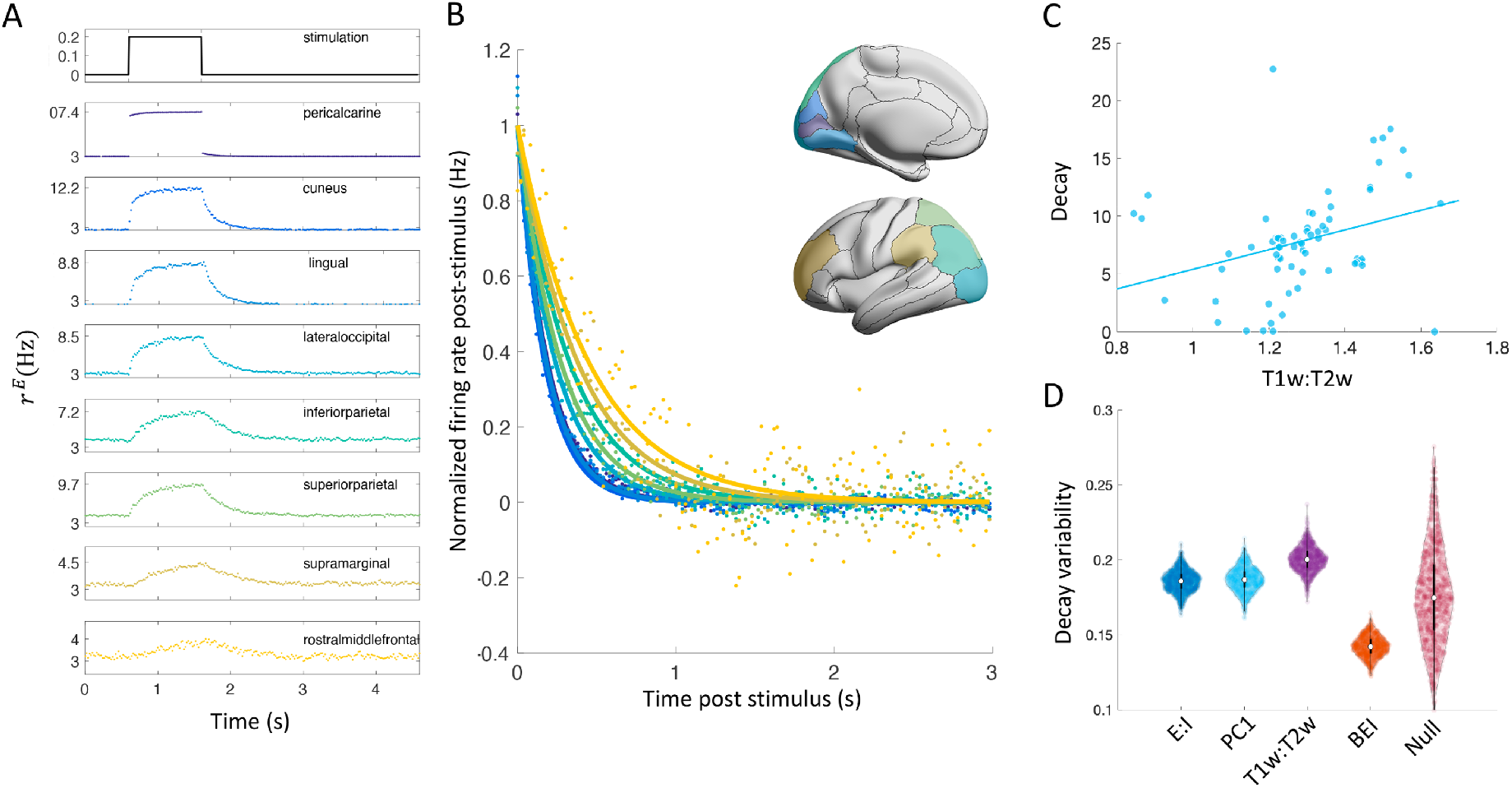
Decay timescales of regional dynamics. **A:** Response time courses for different regions in the E:I model during focal stimulation of pericalcarine cortex. Top plot shows the stimulation time course. The remaining plots show responses in different regions moving through the cortical hierarchy from sensory (near top) to association (near bottom) areas. The *y*-axis in each plot presents the excitatory firing rate (note in pericalcarine the max is 107.4) in Hz, denoted *r*^*E*^. **B:** Decay curves for each of the regions depicted in panel **A**, showing how regional activity returns to baseline following focal stimulation of visual cortex. Surface renderings show the anatomical locations of each region. **C:** Spatial correlation between regional decay rates (in *s*^−1^) and T1w:T2w. Regions with more rapid decay, and thus faster timescales, show higher T1w:T2w. **D:** Distributions of regional variability in decay rates obtained for 1000 runs of each model.

We define the richness of regional timescales in the model as the standard deviation of decay rates across target areas, such that high values reflect a greater diversity of timescales across the brain, which implies a broader dynamic range for information processing. We term this measure regional decay variability. Figure 4D shows that regional decay variability is significantly higher for all biologically constrained heterogeneous models compared to the BEI and spatial null model, with the BEI model showing lower decay variability than the spatial null (all *p*_*bonf*_ < .05). The T1w:T2w model showed greater variability than the E:I and PC1 models, which did not significantly differ from each other (*p*_*bonf*_ = .07). These results suggest that heterogeneity in regional population gain plays an important role in generating a hierarchy of regional timescales. Transcriptionally constrained models show higher variability than spatially rotated null models, but the highest variability is seen in the T1w:T2w model. Thus, variations in myeloarchitecture may help to broaden the dynamic range of information processing beyond regional differences in E:I receptor gene expression.

## Discussion

It has long been recognized that cortical areas vary along multiple dimensions, including cytoarchitecture, myeloarchitecture, chemoarchitecture, and inter-regional connectivity (*8*– *10, 36*). The specific dynamical consequences of each of these variations remain a mystery, but they are likely to have an ultimate influence on the local ratio of excitatory to inhibitory cell activity, resulting in a variable E:I balance across different brain regions. Incorporating such heterogeneity into large-scale models of neuronal dynamics has proven challenging, but the recent availability of diverse data that can be used to index different forms of cellular, molecular, and anatomical variability across brain areas has provided a new means for imposing regional heterogeneity in such models, thus offering an opportunity to understand in detail the dynamical effects of different forms of regional heterogeneity.

To this end, we compared models in which regional heterogeneity was constrained by transcriptional measures of E:I receptor gene expression, the dominant mode of brain-specific gene expression, and T1w:T2w, which is sensitive to variations in myeloarchitecture and which closely tracks regional position in the cortical hierarchy (*27*). We show that the transcriptional E:I model more faithfully reproduces empirical properties of static and dynamic FC while also showing the highest ignition capacity and a broad range of regional activity timescales. Our findings thus highlight a central role for regional variations in E:I balance in generating ignition-like dynamics and timescale hierarchies, and suggest that transcriptional atlas data provide a viable means for constraining heterogeneity in large-scale models of brain activity.

### Evaluating different forms of regional heterogeneity

The starting point for our analysis was an evaluation of how each model captured diverse empirical properties of FC. Early work focused on reproducing the correlation between model and empirical static FC matrices at the edge-level (*11, 30*). Subsequent studies introduced additional constraints, such as evaluating the fit to dynamic FC properties (*25*) or regional variations in average FC (*22*). To our knowledge, no prior study has evaluated all three properties simultaneously. Our analysis shows that edge- and node-level measures of static FC offer loose constraints for model optimization, showing comparably high fit statistics across a broad range of values of the global coupling parameter, *G*. In contrast, fits to FCD show a clear optimum (at *G* = 2.1), mirroring similar results reported previously (*25*). Fitting models to both static and dynamic properties of FC is thus important for identifying an appropriate working point for each model.

Across all three FC properties, heterogeneous models provide a better match to the data than the nominally homogeneous BEI model. This result suggests that regional heterogeneity plays a role in shaping the empirical FC architecture of spontaneous BOLD-dynamics. Critically, the E:I model was the best-performing model across all three benchmarks, indicating that constraining regional heterogeneity by transcriptional markers of E:I balance yields a more faithful replication of empirical FC. However, the differences in fit statistics between models are small. For example, the median static edge-level FC correlation between model and data was r=0.71 for the E:I model, r=0.69 for the T1w:T2w and PC1 models, and r=0.67 for the BEI model. Moreover, the fits to FCD ranged between *D*_*KS*_ = 0.08 (E:I model) and *D*_*KS*_ = 0.1 (BEI model). These results suggest that these empirical fit statistics have only a limited capacity to tease apart dynamical differences between the models.

The model-based evaluations of ignition capacity and decay variability provide a more complete picture. In both cases, the differences between the homogeneous BEI model and the three heterogeneous models were marked: there is a 154% difference in median global ignition capacity values for the E:I and BEI models and a 26% difference in median decay variability values. All heterogeneous models showed notable ignition capacity and decay variability, being higher, on average, than the spatially rotated null model. In contrast, the BEI model showed lower ignition and decay variability than the null model. This result indicates that both ignition and decay depend on some degree of regional heterogeneity, and that biologically plausible forms of heterogeneity increase these properties beyond the expectations of a generic spatial gradient (as represented by the spatially constrained null ensemble).

The E:I model showed the greatest ignition capacity, indicating that regional differences in E:I receptor gene expression may shape local dynamics in a way that promotes rapid and high-fidelity communication. The E:I model also showed a wide range of decay timescales, although the T1w:T2w model showed the highest decay variability, implying that regional variations in myeloarchitecture may additionally enhance the dynamic range of regional processing. Whether increased decay variability in a model provides a more faithful representation of neural dynamics remains unclear. Further work could aim to validate model results against either invasive recordings, which allow precise estimation of decay timescales to focal perturbations, or through specifically designed functional MRI experiments (*35, 37*). Critically, we evaluated the dynamical properties of each model at their empirically optimal working points, meaning that the models show these properties in a parameter regime that closely matches actual data. This is an important constraint, given that prior modelling work evaluating ignition dynamics and timescale hierarchies in Macaque did not fit empirical FC data (*3, 15*). It is thus difficult to ascertain the biological plausibility of the dynamical regime that was studied. The E:I model considered here not only showed the best fit to empirical data, but it also showed the highest ignition capacity and a broad range of decay timescales, suggesting that it offers a parsimonious account of these various dynamical phenomena.

### How does heterogeneity shape neuronal dynamics?

In our models, we introduced heterogeneity by modifying the regional excitability of local population activity. We achieved this by modifying each region’s gain response function. Different approaches have been used to incorporate regional heterogeneity in past work. For example, (*15*) used a hierarchical ordering of cortical areas in macaque based on laminar patterning of inter-regional connectivity to linearly scale the excitatory input strength of each population. In (*22*), T1w:T2w was used to linearly scale the strengths of local excitatory-to-excitatory and inhibitory-to-inhibitory populations when modelling cortical BOLD FC in Human. More complicated approaches are also possible (e.g., (*23*)). We chose to impose heterogeneity by modulating population gain response functions, since local variations in E:I balance will affect the net excitability of the population, which is captured by the gain function parameter, *M*_*i*_. We thus assume that changes in regional gain are the final common effect of different ways in which heterogeneity might influence a specific population or interaction between populations. Critically, regional variations of *M*_*i*_ are modulated by just two terms––the bias, *B*, and the scaling factor, *Z*. This leads to a marked improvement in computational efficiency when compared to more highly parameterized models (e.g., (*22, 23*)), which require the use of heuristic optimization algorithms that can make it difficult to guarantee a globally optimum solution and can shift some parameter values from empirically-constrained ranges. Nonetheless, we expect that further refinement of optimization algorithms capable of robustly and effectively searching high-dimensional parameter spaces will allow greater precision for evaluating different forms of model heterogeneity.

### Implications for understanding ignition-like dynamics

Ignition-like dynamics are thought to be a necessary pre-condition of conscious experience (*4, 33*). As the intensity of a focal sensory stimulus increases, the ignition model predicts that a threshold is reached beyond which widespread activation of other areas is rapidly triggered. Stimuli that do not reach this threshold may only be perceived subconsciously.

Our results indicate that regional heterogeneity is essential for producing ignition-like dynamics, with the E:I model showing the highest ignition capacity. To the extent that our transcriptional E:I measures can be taken as a proxy for receptor abundance (see below), this result suggests that regional variations specifically in E:I receptor activity may play an especially critical role in supporting rapid, high-fidelity broadcasting of sensory stimuli throughout the network.

Recent work using a dynamical model simulated on the Macaque connectome also showed evidence of ignition-like dynamics (*3*). Critically, the authors showed that removal of feedback projections and randomization of the connectivity weights of the SC matrix were each sufficient to abolish this dynamical behaviour, suggesting that feedback connectivity and heterogeneity in SC connection strength give rise to ignition-like activity. In our analysis, the SC matrix is derived from diffusion MRI data and thus cannot distinguish feedforward from feedback connections, although we did incorporate empirical variability in connection strengths. Thus, the fact that we only observe ignition-like dynamics with heterogeneous but not homogeneous models suggests that feedback connectivity is not necessary for ignition, heterogeneity of inter-regional connection weights alone is not sufficient, and that heterogeneity of connection weights and regional population dynamics may represent minimal requirements for the emergence of ignition-like propagation through the network. One caveat to this view is that while the focal perturbation in our analysis did propagate across different levels of the hierarchy, we did not observe widespread activation of frontoparietal systems, which is thought to be a cornerstone of global ignition and which was observed in the macaque model (*3*). This discrepancy could reflect the necessity of feedback connections for generating truly global ignition, or it may reflect a distinct working point of the macaque model, given that it was not constrained by empirical FC data. Further work could specifically try to isolate which model elements represent both necessary and sufficient conditions for cortical ignition.

### Implications for understanding timescale hierarchies

Our analysis indicates that any form of heterogeneity enhances the dynamic range of regional decay timescales compared to the homogeneous BEI model, given that even the spatially rotated null model displayed greater decay variability. Transcriptionally constrained E:I and PC1 models displayed comparable variability, both being greater than the BEI and null models, but the T1w:T2w model showed the highest variability. This result implies that regional myeloarchitecture may play an important role in shaping timescale differences between regions. We note however that part of this additional variability may have been driven by data considerations. We had T1w:T2w measures for both right and left hemispheres, but only gene expression data for the left hemisphere. To generate whole-brain simulations, we thus reflected the left hemisphere expression values to constrain right hemisphere dynamics. Although this may be justified given the previously-noted lack of statistical differences between hemispheres in regional gene expression within the Allen Atlas (*26*), our approach may still have reduced the degree of regional heterogeneity incorporated into the transcriptional models relative to the T1w:T2w model.

Past work has shown that several other properties correlate with various measures of regional activity timescales. In a comprehensive survey of 6390 different regional time series properties estimated for fMRI recordings in the mouse brain, Sethi et al. (*19*) found that timescale-related measures such as relative high-frequency signal power, correlate with structural measures of in-degree, such that the time series of regions with many incoming connections showed greater low-frequency power. Similar results were later reported in human (*18*). Other work in humans has also shown that regional variations in autocorrelation and other time series properties track the cortical hierarchy (*20, 38*). These additional measures could be used to provide additional constraints on model fitting, although the temporal resolution of functional fMRI may be too limited to provide a detailed understanding of regional variations in activity timescales.

### Limitations

We relied on a single coarse scale parcellation of the brain to allow exhaustive parameter sweeps over several different models. This choice was essential to reduce computational burden, and it ensures that our results are directly comparable with other studies using the same parcellation (e.g., (*11*)). Nonetheless, parcellation type and scale can have an effect on empirical connectome properties (*39, 40*), and further evaluation of the degree to which our results generalize to other parcellations is required.

Our E:I model used regional measures of excitatory and inhibitory receptor gene expression as a proxy for the relative abundance of each receptor type in each region, which in turn is assumed to be related to the relative activity levels of inhibitory and excitatory cells. Gene expression in the AHBA was assayed using microarray, which quantifies messenger RNA (mRNA) abundance. The correspondence between mRNA levels and protein abundance can be complex and may vary across different tissues or brain regions (*41*). Moreover, the precise correspondence between receptor densities and cell-specific activity levels remains unclear. The increasing availability of single-cell RNAseq data will provide more precise estimates of regional cell-type abundances (*42*). Any imprecision in our estimates should limit the ability of the E:I model to yield physiologically-plausible dynamics. Our results may therefore represent a lower bound on the accuracy that such a model may achieve.

### Conclusions

We show that regional heterogeneity improves model fits to empirical FC properties, particularly when models are constrained by transcriptomic estimates of E:I receptor gene expression. Critically, at the same working point, this model also shows the highest ignition capacity and a broad range of regional activity timescales, suggesting that regional variations in the activity levels of excitatory and inhibitory receptors may play an important role in the emergence of these dynamical properties. Our analysis indicates that recently constructed transcriptional atlas data provide a fruitful and biologically principled means for imposing regional heterogeneity in large-scale dynamical models.

## Methods

### Empirical MRI data

#### Image acquisition

Eyes-closed resting-state fMRI data were acquired in 389 healthy individuals (170 males; mean age = 24 years; SD = 4.2 years) using a Siemens Skyra 3 Tesla scanner with a 32-channel head coil located Monash Biomedical Imaging in Clayton, Victoria, Australia using the following parameters: time-to-repetition (TR), 754 ms; time-to-echo (TE), 21 ms; flip angle 50^º^; multiband acceleration factor, 3; field-of-view, 190 mm; voxel size, 3 mm^3^ isotropic. A total of 620 volumes were acquired. T1-weighted structural images were also acquired using 1mm^3^ isotropic voxels; TR = 2300 ms; TE = 2.07 ms; FOV=256 ⨯ 256 mm. Diffusion MRI data were available for a subset of 289 individuals, acquired using the following parameters: 2.5mm3 voxel size, TR = 8800 ms, TE = 110 ms, FOV of 240 × 240 mm, 60 directions with b = 3000 s/mm2 and seven b = 0 volumes. A single b = 0 s/mm2 volume was obtained with the reverse-phase encoding for use in distortion correction.

#### DWI processing and Connectome construction

The diffusion data were processed using MRtrix3 version 3.0 (https://www.mrtrix.org/) and FSL version 5.0.11 (https://fsl.fmrib.ox.ac.uk/fsl/fslwiki), as detailed in Oldham et al. (*43*). Briefly, the diffusion images for each individual were corrected for eddy-induced current distortions, susceptibility-induced distortions, inter-volume head motion, outliers in the diffusion signal (*44*), within-volume motion (*45*), and B1 field inhomogeneities (*46*). Tractography was conducted using the Fibre Orientation Distributions (iFOD2) algorithm, as implemented in MRtrix3 (*47*), which utilises fibre orientation distributions estimated for each voxel using constrained spherical deconvolution, which can improve the reconstruction of tracts in highly curved and crossing fibre regions (*48, 49*). Streamline seeds were preferentially selected from areas where streamline density was under-estimated with respect to fibre density estimates from the diffusion model (*50*).

We used Anatomically Constrained Tractography to further improve the biological accuracy of streamlines (*51*). To create a structural connectivity matrix, streamlines were assigned to each of the closest regions in the parcellation within a 5mm radius of the streamline endpoints (*52*), yielding an undirected 82×82 connectivity matrix. A group-representative connectome was generated by selecting 30% of edges showing the lowest coefficient of variation (based on the streamline count) across participants (*53*).

#### fMRI processing and FC estimation

The functional MRI data were processed as detailed in (*54*). Briefly, the functional images were initially preprocessed in FSL FEAT following a basic pipeline, which included removal of the first four volumes, rigid-body head motion correction, 3mm spatial smoothing to improve signal-to-noise ratio, and high pass temporal filter of 75s to remove slow drifts (*55*). The data were then denoised using FSL-FIX, an independent component analysis (ICA)-based denoising method that uses an automated classifier to identify noise-related components for removal from the data (*56*). The algorithm was trained on a manually labelled held-out set of 25 individuals scanned with identical imaging parameters. Time courses for noise-labelled components, along with 24 head motion parameters (6 rigid-body parameters, their backwards derivatives, and squared values of the 12 regressors), were then removed from the voxelwise functional MRI time series using ordinary least squares regression.

Denoised functional data were spatially normalized to the International Consortium for Brain Mapping 152 template in Montreal Neurological Institute (MNI) space using ANTs (version 2.2.0) (*57*), via a three-step method: 1) registration of the mean realigned functional scan to the skull-stripped high resolution anatomical scan via rigid-body registration; 2) spatial normalization of the anatomical scan to the MNI template via a nonlinear registration; and 3) normalization of functional scan to the MNI template using a single transformation matrix that concatenates the transforms generated in steps 1 and 2. Mean time series for each parcellated region were then extracted and Pearson correlations between each pair of regional time series were used to estimate inter-regional FC matrices. Functional connectivity dynamics (FCD) were also calculated for the empirical data, as outlined below.

### The Balanced Excitation-Inhibition (BEI) Dynamic Mean Field Model

We simulated spontaneous neuronal activity using a whole-brain model that couples the local population dynamics of different brain areas through an anatomical inter-regional connectivity matrix derived from diffusion-weighted MRI data. The local population dynamics of each area are described by a dynamic mean field model, as proposed by (*11*). The model expresses the activity of large ensembles of interconnected excitatory and inhibitory spiking neurons as a reduced set of dynamical equations describing the population rate activity of coupled excitatory (E) and inhibitory (I) pools of neurons, following the original derivation of Wong and Wang (*58*). At the local neuronal population level, the excitatory synaptic currents, *I*^*(E)*^, are mediated by NMDA receptors and the inhibitory currents, *I*^*(I)*^, are mediated by GABA_A_ receptors. Within each brain area, the E and I neuronal pools are mutually connected. The inter-area coupling between local neuronal population dynamics corresponding to two different brain areas *i* and *j* is mediated only at the E-to-E level and defined by the scaled structural connectivity matrix *C*. The elements of this matrix, *C*_*ij*,_ define the inter-regional anatomical connectivity with connection weights estimated using tractography of diffusion-weighted MRI data, as described above.

The balanced dynamic mean-field whole-brain model is mathematically described by the following system of coupled differential equations:

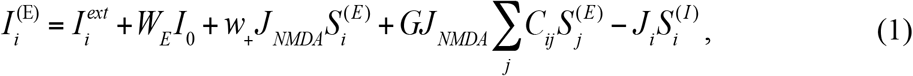

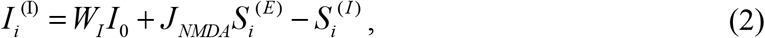

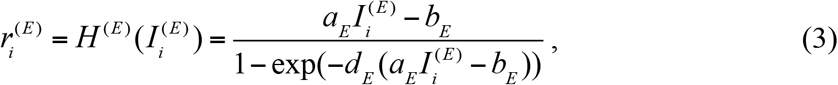

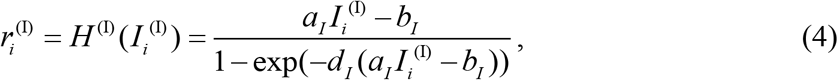

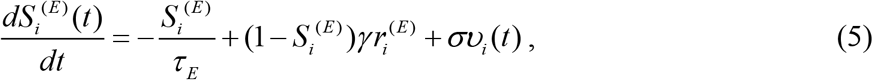

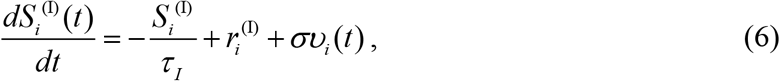

In the last equations, for each excitatory *(E)* or inhibitory *(I)* pool of neurons in each brain area *i, r*_*i*_^*(E,I)*^ (in Hz) indicates the firing rate, *I*_*i*_^*(E,I)*^ (in nA) denotes the total input current, and *S*_*i*_^*(E,I)*^ represents the synaptic gating variable. The total input current received by the E and I pools is transformed into firing rates, *r*_*i*_^*(E,I)*^, by *H*^*(E,I)*^, which corresponds to the sigmoidal neuronal response functions defined in (*59*). In the excitatory *(E)* and inhibitory *(I)* neuronal response functions, *g*_*E*_ = 310 nC^-1^ and *g*_*I*_ = 615 nC^-1^ define the gain factors determining the slope of *H, I*_*thr*_^*(E)*^ = 0.403 nA, and *I*_*thr*_^*(I)*^= 0.288 nA are the threshold currents above which the firing rates increase linearly with the input currents, and *d*_*E*_ =0.16 and *d*_*I*_ =0.087 are constants determining the shape of the curvature of *H* around *I*_*thr*_, respectively. The synaptic gating variable of excitatory pools, *S*_*i*_^*(E)*^, is mediated by NMDA receptors with a decay time constant *τ*_*NMDA*_ = 0.1 s and *γ* = 0.641, whereas the average synaptic gating in inhibitory pools is mediated by GABA receptors with a decay time constant *τ*_*GABA*_ = 0.01s. The local external input impinging in each population is *I*_*0*_ = 0.382 nA and is weighted with *W*_*E*_ =1 for the excitatory pools and with *W*_*I*_=0.7 for the inhibitory pools. The parameter 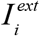captures external stimulation to the excitatory population. In all cases, we set 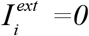 for all regions, except when assessing the ignition capacity and response decay profiles of the model, in which case we set 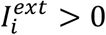, as detailed below. The local recurrent excitation in each excitatory pool is weighted by *w*_*+*_=1.4. All excitatory synaptic couplings are weighted by *J*_*NMDA*_=0.15 nA. In equations (5) and (6) *ν*_*n*_ denotes uncorrelated standard Gaussian noise with an amplitude of *σ*=0.01 nA. All parameters were set as in (*11*).

One method for introducing heterogeneity to this model is to define the parameters such that each isolated brain area shows asynchronous spontaneous activity with low firing rate (*r*^*(E)*^∼3Hz and *r* ^*(I)*^∼9Hz), as shown in in electrophysiological experiments (*12*–*14*). In this regime, the whole-brain model is balanced. The balance is obtained by adjusting each local feedback inhibition weight, *J*_*i*_, for each brain area *i* such that the firing rate of the excitatory pools *r*_*i*_^*(E)*^ remains clamped at 3Hz even when receiving excitatory input from connected areas. This regulation for achieving balance is known as Feedback Inhibition Control (FIC) and is obtained numerically by the method described in (*11*). Indeed, this work demonstrated that FIC improves the fitting of empirical resting-state FC and more crucially, achieves more realistic evoked activity. We refer to this model as the balanced excitation-inhibition (BEI) model. Despite the regional heterogeneity introduced by local tuning of the FIC parameter, we use this model as our homogeneous benchmark for comparing to other models, since the FIC adjustments ensure uniform firing rates across all regions and all regional gain parameters are fixed (as detailed below).

The BEI model is fitted to empirical fMRI data by finding the optimal global coupling factor *G*, which uniformly scales all inter-area E-to-E connections. The parameter *G* is the only control parameter that is adjusted to move the homogeneous model to its optimal working point, where the simulated activity maximally fits the empirically-observed resting-state FC architecture. In particular, we fit static edge-level FC, node-level FC, and FCD, by choosing the coupling factor *G* such that the correlation between the empirical and simulated FC is high and the distance between the data and model FCD distributions is minimal (i.e., the minimal *D*_*KS*_ between empirical and simulated FCD; see below).

Simulations were run for a range of *G* between 0 and 3 with an increment of 0.01 and with a time step of 0.1 ms. For each value of *G*, we averaged 389 simulations of 464.464 seconds each, which were then put through a BOLD forward model (see below) in order to emulate the empirical resting-state data, which comprised haemodynamic recordings in 616 volumes acquired in each of 389 human subjects. We optimized the coupling factor *G* before introducing regional heterogeneity to the model using the gain and scale parameters (i.e., B=0 and *Z*=0 during this fitting procedure; see below).

### Simulating BOLD signals

Regional BOLD signals are simulated in the model using the generalized hemodynamic model of (*60*). We calculate the BOLD signal in each brain area *i* from the simulated firing rate of the excitatory pools *r*_*i*_^*(E)*^. In this hemodynamic model, an increase in the firing rate causes an increase in a vasodilatory signal, *s*_*i*,_ that is subject to auto-regulatory feedback. Blood inflow, *f*_*i*_, responds in proportion to this signal, inducing changes in blood volume, *v*_*i*_, and deoxyhemoglobin content, *q*_*i*_. The differential equations describing the hemodynamic model coupling these biophysical variables are:

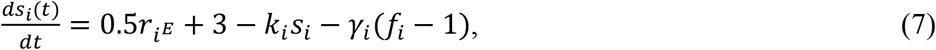

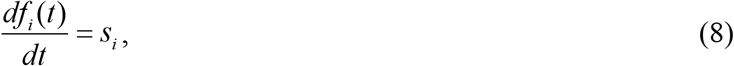

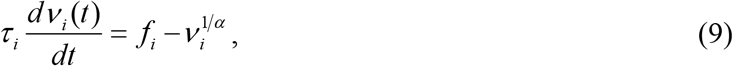

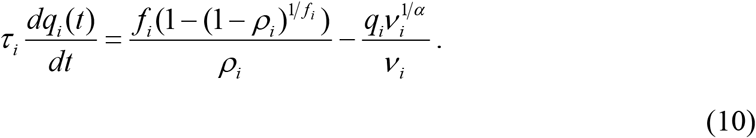

In these equations, *ρ* denotes the resting oxygen extraction fraction, τ is a time constant and α represents the resistance of the veins. In order to compute in each area *i*, the BOLD signal, *B*_*i*_, we calculate a volume-weighted sum of extra- and intravascular signals, which comprises a static nonlinear function of volume, *v*_*i*_, and deoxyhemoglobin content, *qi*, and is expressed as follows:

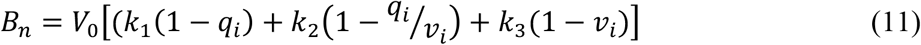

We adopted the biophysical parameters recommended by (*60*). To focus on the most functionally relevant frequency range in resting-state conditions, we bandpass filtered both empirical and simulated BOLD signals (0.08 > *f* > 0.008 Hz).

### Introducing regional heterogeneity

One of the key drivers of local population activity in the dynamical model is the balance between excitatory and inhibitory activity. E:I balances across brain regions are thought to play a critical role in synchronized dynamics and their disruptions have been implicated in diverse diseases (*61, 62*). In the absence of accurate density estimates for each cell type in different regions of the human brain, regional variations in E:I balance could, in principle, be parameterized in many different ways. Here, we use data on regional expression levels of excitatory and inhibitory receptors to constrain our model of E:I heterogeneity across the brain. To this end, we use data from the Allen Institute Human Brain Atlas (AHBA), which comprises microarray data quantifying the transcriptional activity of >20,000 genes in >4,000 different tissue samples distributed throughout the brain, taken from six post-mortem samples (*26, 29*).

The AHBA data were processed according to the pipeline developed in Arnatkevicuite et al. (*28*). Briefly, probe-to-gene annotations were updated using the Re-Annotator toolbox (*63*), resulting in the selection of 45,821 probes corresponding to a total of 20,232 genes. Tissue samples derived from brainstem and cerebellum were removed. Then, probes that did not exceed background noise in more than 50% of remining samples were removed, excluding 13,844 probes corresponding to 4,486 genes. A representative probe for each gene was selected based on the highest mean intensity across regions (*64*). To increase the anatomical accuracy of sample-to-region matching, samples were first divided into four separate groups based on their location: hemisphere (left/right) and structure assignment (cortex/subcortex) and then assigned to the corresponding parcellation-defined regions of interest separately for each of the four groups. Donor-specific grey matter parcellations were generated and samples located within 2 mm of the parcellation voxels were mapped to the closest voxel assigning almost 90% of all tissue samples. We retained only data for the left cortex, as anatomical coverage in the AHBA is more complete for this hemisphere. Finally, gene-expression measures within a given brain were normalized using a scaled robust sigmoid normalization for every gene across samples. Normalized expression measures in samples assigned to the same region were averaged and aggregated into a region × gene matrix consisting of expression measures for 15,745 genes over 34 left hemisphere regions. Given the lack of notable inter-hemispheric differences in regional gene expression identified in the AHBA (*26*), the left hemisphere values were reflected on to the right hemisphere to enable whole-brain model simulations.

To capture regional heterogeneity in E:I balance, we extracted expression measures specifically for genes coding for the excitatory AMPA and NMDA receptors and inhibitory GABA-A receptor isoforms and subunits. More specifically, the genes selected for AMPA were GRIA1, GRIA2, GRIA3, GRIA4; the genes for NMDA were GRIN1, GRIN2A, GRIN2B, GRIN2C; and the genes for GABA-A were GABRA1, GABRA2, GABRA3, GABRA4, GABRA5, GABRB1, GABRB2, GABRB3, GABRG1, GABRG2, GABRG3. In order to increase the accuracy of gene expression measures, probes representing the expression of individual genes were selected based on the highest correlation to RNA sequencing data in two of the six donor brains (*64*). In our implementation, we quantify regional E:I balance simply by summing all expression values for AMPA and NMDA, summing all values for GABA, and dividing the former by the latter, resulting in an E:I ratio normalized to the [0,1] interval. For each brain region *i*, we denote this normalized gene expression-based excitation/inhibition ratio by *R*_*i*_. We note that estimating E:I balance from gene expression data is challenging given that each receptor is a heteromer, consisting of differing subunit compositions (AMPAR: heterotetrameric assembly from 4 different subunits; NMDAR: heterotetrameric assembly from 7 different subunits; GABA-AR: homo- and heteropentameric assembly from 19 subunits)(*65*–*68*). The exact combination of these subunits can alter receptor properties, including biophysical, pharmacological and signalling attributes. For example, the GluN2 subunit is essential for determining the gating properties of NMDA receptors, with different subunit compositions impacting excitatory postsynaptic decay times (*68*). Here we have adopted a pragmatic approach and chosen genes encoding the major subunit types present in the human adult cortex for each receptor, assuming the relative expression levels provide an adequate estimate of the E:I ratio within a given cortical region.

We used the transcriptomically-constrained local E:I ratio *R*_*i*_ to modulate the gain of the local neuronal response functions of the corresponding excitatory and inhibitory pools of each brain region, *i*, following a number of experimental and theoretical studies (*69, 70*). Specifically, we consider that *R*_*i*_ modulates the gain of the neuronal response function *H*^*(E,I)*^ in each brain area according to the modified equations (3) and (4):

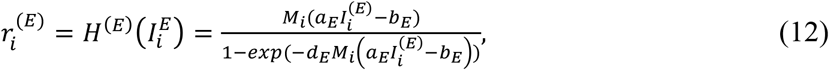

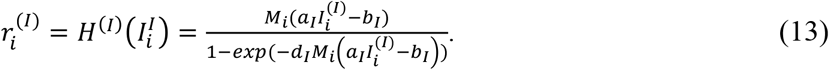

Similar to (*22*), we assume that the heterogeneity factors (in our case, the transcriptomically-defined local excitation/inhibition ratios *R*_*i*_), are linearly transformed by two unknown free parameters corresponding to a bias *B* and a scaling factor *Z*. Thus, in equations (12) and (13), *M*_*i*_ = 1 + *B* + *ZR*_*i*_. In other words, the effect of the transcriptomically-constrained E:I ratio *R*_*i*_ is to modulate locally the slope of the corresponding excitatory and inhibitory neuronal response functions, *H*. We studied exhaustively the two free parameters *B* and *Z* regulating the influence of transcriptional heterogeneity. Homogeneous conditions correspond to cases where *Z=0*. Note that the balancing FIC procedure is applied after fixing the gain scaling, i.e. for a specific bias *B* and a specific scaling factor *Z*. Also note that we although constrained the model using a transcriptional index of E:I balance, we used this constraint to modify the gain of each population model. We did this under the assumption that the population gain function, capturing net regional excitability, will be sensitive to regional variations in E:I balance, and in order to limit the number of fitted parameters in the model. More specifically, we fitted only two parameters—*B* and *Z*—to model regional heterogeneity whereas other approaches have fitted four or more (*22, 23*). We adopted this strategy to enable a more comprehensive search of parameter space, which is important to identify a unified working point for all outcome properties of the model. A more direct modulation of E:I weights might yield different results.

We additionally evaluated two additional forms of biologically-constrained heterogeneity: the PC1 and T1w:T2w models. The PC1 model was also informed by AHBA data. In this case, we identified the dominant mode of spatial variation in gene expression as the first principal component (PC1) of 1,926 genes that overlapped between our final quality-controlled gene list and those previously deemed to be brain-specific in (*27*), where it was shown that this mode correlates with T1w:T2w.

The T1w:T2w model incorporated heterogeneity by directly relying on empirical estimates of the T1w:T2w ratio, which is sensitive to myelin content (*71*). Prior work has shown that this measure can also be used to map cortical hierarchies (*27*), and that large-scale heterogeneous biophysical models constrained by T1w:T2w more accurately reproduce empirical FC properties than homogeneous models (*22*). Regional T1w:T2w values were obtained from the Human Connectome Project dataset, averaged over 1200 subjects following 2 mm FHWM Gaussian smoothing in grayordinates space (https://balsa.wustl.edu/study/show/RVVG), and then parcellated using the Desikan-Killany atlas (*34*).

Finally, to further ensure that any results obtained for the E:I model cannot be explained by a generic spatial gradient, we also evaluated model performance after permuting the assignment of E:I values to regions. A total of 10,000 permutations were performed by spatially rotating the expression values using the approach described in (*72*). Critically, this approach preserves the spatial autocorrelation of the expression values, which is essential for ensuring that any resulting effects are not driven by low-order spatial gradients (see also (*31*)). In all cases, we normalized the corresponding heterogeneity factor to the [0,1] interval, and treat it similarly to the ratio *R*_*i*_.

### Evaluating model performance

Models were fitted to three empirical properties of the FC data, as detailed in the following. To compare model performance, we simulated 1000 runs of each model at its own optimal parameters to obtain a distribution of fit statistics that captures the inherent stochasticity of the model.

#### Static edge-level FC

The static edge-level FC is defined as the *N*×*N* matrix of BOLD signal correlations between brain areas computed over the entire recording period (see Figure 1B). We computed the empirical FC for each human participant and for each simulated trial (the total number of trials matched the number of participants). The group-averaged empirical and simulated FC matrices were compared by computing the Pearson correlation between their upper triangular elements (given that the FC matrices are symmetric).

#### Static node-level FC

Node-level FC, or node strength, is another static spatial measure that characterizes the average FC strength for each area (see Figure 1E) (*73*). It has also been called global brain connectivity in previous work (*22*). Thus, node-level functional connectivity strength is defined as

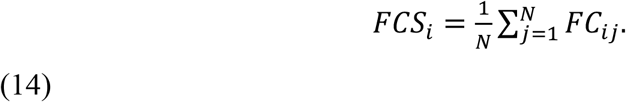

We computed the empirical strength vectors for each human participant and for each simulated trial (total number of trials matched the number of participants). The node FC fit is quantified by computing the Pearson correlation between the group averaged empirical and simulated strength vectors.

#### Functional Connectivity Dynamics (FCD)

The spatiotemporal dynamics of resting-state activity are captured by the statistics of how FC computed over sliding windows evolves across time (see Figure 1E). Here, we computed the FC over a sliding window of 80 TRs (corresponding approximately to 1 minute) with incremental shifts of 18 TRs. In order to quantify the spatiotemporal statistics of time-evolving FC, we calculated a time-versus-time matrix, called Functional Connectivity Dynamics (*FCD*) (*25*). This FCD matrix is defined so that each entry, *FCD*(*t*_*x*_, *t*_*y*_), corresponds to the correlation between the *FC* centred at times *t*_*x*_ and the *FC* centred at *t*_*y*_. The statistical distribution of *FCD* values captures the spatiotemporal architecture of the recording session. In order to compare quantitatively the spatiotemporal dynamical characteristics between empirical data and model simulations, we generate the distributions of the upper triangular elements of the FCD matrices over all participants as well as of the FCD matrices obtained from all simulated trials for a given parameter setting. The similarity between the empirical and model FCD distributions is then compared using the Kolmogorov-Smirnov distance, *D*_*KS*_, allowing for a meaningful evaluation of model performance in predicting the changes observed in dynamic resting-state FC.

#### Ignition capacity

The ignition capacity of each model was quantified by examining how regional dynamics respond to a simulated focal perturbation. First, for a given working point, we run the whole-brain model for 3000 ms under spontaneous conditions in order to eliminate transients. Second, from 3000 to 4000 ms, we apply a stimulation of strength 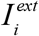to one of four occipital regions, *i*, bilaterally in both hemispheres (these regions were the lateral occipital, pericalcarine, cuneus, and lingual brain areas in the Desikan-Killiany atlas). Third, from 4000 ms to 7000 ms we reset the external stimulation to zero again, and let the system relax again to the spontaneous state. For each condition, i.e., for each parameter setting, and each particular stimulation 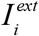, we run 30 trials and compute the average, across trials, of the elicited activity as a function of time on all brain areas, in order to smooth the resulting neuronal whole-brain activity.

To compare ignition capacity across the whole cortex, we fix the location *i* where the stimulation is applied (and also stimulate the region’s contralateral homologue). Then, we simulate the averaged neuronal response elicited in a given brain area as a consequence of the applied artificial stimulation and as a function of the strength 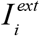 (we modify 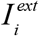 from 0 to 0.2 in steps of 0.001). For each brain region, we perform a sigmoidal fitting of the elicited averaged rate activity in the period from 3500 to 4000 ms as a function of the applied strength 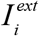 in the stimulated specific site *i*. The ignition elicited for a given target brain area after the application of the artificial stimulation in the specific region *i* is defined by the product between the level of concavity of the neuronal response function elicited on that target area and the maximal rate elicited when the maximal strength of stimulation is applied (as fitted by the sigmoidal function). The level of concavity is defined by the maximal value of the second derivative of the fitted sigmoidal function. Figure 1F-G shows graphically the key concept. For a given stimulation site *i*, the global ignition is the average ignition across all brain areas. Finally, the ignition is defined by averaging the global ignition over all stimulated sites *i*. We stimulated lower-order occipital regions to study the capacity of the system to propagate a sensory perturbation to the rest of the brain (*3*). The ignition measure is basically describing two effects, namely the fidelity (captured by the maximal elicited rate in the target area) and explosiveness (captured by the level of concavity) of ignition-like activity.

#### Decay timescales

Regional decay timescales were estimated using the same stimulation paradigm as used for quantifying ignition. More specifically, we compute the temporal decay of each target brain region following simulation of the pericalcarine region in both hemispheres. The decay rate was calculated by fitting an exponential curve to the regional firing rates post-stimulation, averaged over 1000 simulations (the length of total simulation as described above). The functional form of the exponential curve was 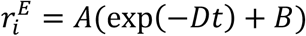, where 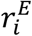 is the firing rate (Hz) post-stimulation, *t* is the time post-stimulation (in seconds) and the free parameters of decay rate, *D*, scaling constant, *A*, and offset, *B*, were fitted using nonlinear least squares (implemented in Matlab). We characterized time hierarchy, following (*15*), as the standard deviation of the decays across brain areas (see Figure 1G). A large standard deviation implies greater variability in intrinsic timescales across brain regions, which is consistent with a broader dynamic range of activity and a more pronounced hierarchical ordering of brain areas. The final decays are computed, as before, by averaging over all site stimulations. Note that due to the large number of model iterations in the heterogeneity analysis, the time decay was approximated using a linear fit of the signal decay in log-space for computational efficiency.

### Statistical analyses

Differences in model fits to empirical properties, as well as in ignition capacity and decay variability, were assessed using pair-wise Wilcoxon ranksum tests, Bonferroni-corrected for 10 possible comparisons (corrected *p*-values are denoted *p*_*bonf*_). Spatial correlations between regional maps were assessed with reference to empirical null distributions generated by rotating one of the spatial maps using the same procedure as used when generating the spatial null models (74). The corresponding *p*-values are denoted *p*_*spatial*_.

## Acknowledgements

G.D. is supported by the Spanish Research Project COBRAS PSI2016-75688-P (AEI/FEDER, EU), by the European Union’s Horizon 2020 Research and Innovation Programme under grant agreements n. 720270 (HBP SGA1) and n. 785907 (HBP SGA2), and by the Catalan AGAUR Programme 2017 SGR 1545. AF was supported by the Sylvia and Charles Viertel Charitable Foundation and National Health and Medical Research Council (ID: 3251549).

## Code and Data availability

The Matlab code and data used in this study and carefully described above are available from the corresponding authors upon reasonable request.

## References

1. E. Bullmore, O. Sporns, Complex brain networks: graph theoretical analysis of structural and functional systems. Nat Rev Neurosci. 10, 186–198 (2009).

2. G. Deco, V. K. Jirsa, A. R. McIntosh, Emerging concepts for the dynamical organization of resting-state activity in the brain. Nat Rev Neurosci. 12, 43–56 (2011).

3. M. R. Joglekar, J. F. Mejias, G. R. Yang, X.-J. Wang, Inter-areal balanced amplification enhances signal propagation in a large-scale circuit model of the primate cortex. Neuron. 98, 222–234.e8 (2018).

4. G. A. Mashour, P. Roelfsema, J.-P. Changeux, S. Dehaene, Conscious Processing and the Global Neuronal Workspace Hypothesis. Neuron. 105, 776–798 (2020).

5. G. Deco, V. Jirsa, A. R. McIntosh, O. Sporns, R. Kotter, Key role of coupling, delay, and noise in resting brain fluctuations. Proceedings of the National Academy of Sciences. 106, 10302–10307 (2009).

6. C. J. Honey, R. Kötter, M. Breakspear, O. Sporns, Network structure of cerebral cortex shapes functional connectivity on multiple time scales. Proc Natl Acad Sci U S A. 104, 10240–10245 (2007).

7. G. Deco, V. K. Jirsa, Ongoing Cortical Activity at Rest: Criticality, Multistability, and Ghost Attractors. Journal of Neuroscience. 32, 3366–3375 (2012).

8. K. Brodmann, Vergleichende Lokalisationslehre der Grosshirnrinde (Johann Ambrosius Barth, Leipzig, 1909).

9. X.-J. Wang, Macroscopic gradients of synaptic excitation and inhibition in the neocortex. Nature Reviews. Neuroscience, 1–10 (2020).

10. J. M. Huntenburg, P.-L. Bazin, D. S. Margulies, Large-scale gradients in human cortical organization. Trends in Cognitive Sciences. 22, 21–31 (2018).

11. G. Deco, A. Ponce-Alvarez, P. Hagmann, G. L. Romani, D. Mantini, M. Corbetta, How Local Excitation-Inhibition Ratio Impacts the Whole Brain Dynamics. Journal of Neuroscience. 34, 7886–7898 (2014).

12. B. D. Burns, A. C. Webb, The spontaneous activity of neurones in the cat’s cerebral cortex. Proc R Soc Lond B Biol Sci. 194, 211–223 (1976).

13. K. W. Koch, J. M. Fuster, Unit activity in monkey parietal cortex related to haptic perception and temporary memory. Exp Brain Res. 76, 292–306 (1989).

14. W. R. Softky, C. Koch, The highly irregular firing of cortical cells is inconsistent with temporal integration of random EPSPs. J Neurosci. 13, 334–350 (1993).

15. R. Chaudhuri, K. Knoblauch, M.-A. Gariel, H. Kennedy, X.-J. Wang, A Large-Scale Circuit Mechanism for Hierarchical Dynamical Processing in the Primate Cortex. Neuron. 88, 419–431 (2015).

16. P. Barone, A. Batardiere, K. Knoblauch, H. Kennedy, Laminar Distribution of Neurons in Extrastriate Areas Projecting to Visual Areas V1 and V4 Correlates with the Hierarchical Rank and Indicates the Operation of a Distance Rule. J. Neurosci. 20, 3263– 3281 (2000).

17. J. D. Murray, A. Bernacchia, D. J. Freedman, R. Romo, J. D. Wallis, X. Cai, C. Padoa-Schioppa, T. Pasternak, H. Seo, D. Lee, X.-J. Wang, A hierarchy of intrinsic timescales across primate cortex. Nat Neurosci. 17, 1661–1663 (2014).

18. J. Fallon, P. G. D. Ward, L. Parkes, S. Oldham, A. Arnatkevičiūtė, A. Fornito, B. D. Fulcher, Timescales of spontaneous fMRI fluctuations relate to structural connectivity in the brain. Network Neuroscience. 4, 788–806 (2020).

19. S. S. Sethi, V. Zerbi, N. Wenderoth, A. Fornito, B. D. Fulcher, Structural connectome topology relates to regional BOLD signal dynamics in the mouse brain. Chaos. 27, 047405 (2017).

20. R. V. Raut, A. Z. Snyder, M. E. Raichle, Hierarchical dynamics as a macroscopic organizing principle of the human brain. Proc Natl Acad Sci USA, 202003383 (2020).

21. G. J. Yang, J. D. Murray, X.-J. Wang, D. C. Glahn, G. D. Pearlson, G. Repovs, J. H. Krystal, A. Anticevic, Functional hierarchy underlies preferential connectivity disturbances in schizophrenia. Proc Natl Acad Sci USA. 113, E219–E228 (2016).

22. M. Demirtaş, J. B. Burt, M. Helmer, J. L. Ji, B. D. Adkinson, M. F. Glasser, D. C. Van Essen, S. N. Sotiropoulos, A. Anticevic, J. D. Murray, Hierarchical heterogeneity across human cortex shapes large-scale neural dynamics. Neuron, 1–28 (2019).

23. P. Wang, R. Kong, X. Kong, R. Liegeois, C. Orban, G. Deco, M. P. van den Heuvel, B. T. Thomas Yeo, Inversion of a large-scale circuit model reveals a cortical hierarchy in the dynamic resting human brain. Science advances. 5, eaat7854 (2019).

24. D. S. Margulies, S. S. Ghosh, A. Goulas, M. Falkiewicz, J. M. Huntenburg, G. Langs, G. Bezgin, S. B. Eickhoff, F. X. Castellanos, M. Petrides, E. Jefferies, J. Smallwood, Situating the default-mode network along a principal gradient of macroscale cortical organization. Proceedings of the National Academy of Sciences. 113, 12574–12579 (2016).

25. G. Deco, M. L. Kringelbach, V. K. Jirsa, P. Ritter, The dynamics of resting fluctuations in the brain: metastability and its dynamical cortical core. Sci Rep. 7, 3095 (2017).

26. M. J. Hawrylycz, E. S. Lein, A. L. Guillozet-Bongaarts, E. H. Shen, L. Ng, J. A. Miller, L. N. van de Lagemaat, K. A. Smith, A. Ebbert, Z. L. Riley, C. Abajian, C. F. Beckmann, A. Bernard, D. Bertagnolli, A. F. Boe, P. M. Cartagena, M. M. Chakravarty, M. Chapin, J. Chong, R. A. Dalley, B. D. Daly, C. Dang, S. Datta, N. Dee, T. A. Dolbeare, V. Faber, D. Feng, D. R. Fowler, J. Goldy, B. W. Gregor, Z. Haradon, D. R. Haynor, J. G. Hohmann, S. Horvath, R. E. Howard, A. Jeromin, J. M. Jochim, M. Kinnunen, C. Lau, E. T. Lazarz, C. Lee, T. A. Lemon, L. Li, Y. Li, J. A. Morris, C. C. Overly, P. D. Parker, S. E. Parry, M. Reding, J. J. Royall, J. Schulkin, P. A. Sequeira, C. R. Slaughterbeck, S. C. Smith, A. J. Sodt, S. M. Sunkin, B. E. Swanson, M. P. Vawter, D. Williams, P. Wohnoutka, H. R. Zielke, D. H. Geschwind, P. R. Hof, S. M. Smith, C. Koch, S. G. N. Grant, A. R. Jones, An anatomically comprehensive atlas of the adult human brain transcriptome. Nature. 489, 391–399 (2012).

27. J. B. Burt, M. Demirtaş, W. J. Eckner, N. M. Navejar, J. L. Ji, W. J. Martin, A. Bernacchia Anticevic, J. D. Murray, Hierarchy of transcriptomic specialization across human cortex captured by structural neuroimaging topography. Nature Neuroscience. 21, 1251– 1259 (2018).

28. A. Arnatkeviciute, B. D. Fulcher, A. Fornito, A practical guide to linking brain-wide gene expression and neuroimaging data. Neuroimage. 189, 353–367 (2019).

29. A. Fornito, A. Arnatkevičiūtė, B. D. Fulcher, Bridging the Gap between Connectome and Transcriptome. Trends Cogn Sci. 23, 34–50 (2019).

30. G. Deco, A. Ponce-Alvarez, D. Mantini, G. L. Romani, P. Hagmann, M. Corbetta, Resting-State Functional Connectivity Emerges from Structurally and Dynamically Shaped Slow Linear Fluctuations. Journal of Neuroscience. 33, 11239–11252 (2013).

31. A. F. Alexander-Bloch, H. Shou, S. Liu, T. D. Satterthwaite, D. C. Glahn, R. T. Shinohara, S. N. Vandekar, A. Raznahan, On testing for spatial correspondence between maps of human brain structure and function. Neuroimage. 178, 540–551 (2018).

32. F. Váša, J. Seidlitz, R. Romero-Garcia, K. J. Whitaker, G. Rosenthal, P. E. Vértes, M. Shinn, A. Alexander-Bloch, P. Fonagy, R. J. Dolan, P. B. Jones, I. M. Goodyer, NSPN consortium, O. Sporns, E. T. Bullmore, Adolescent Tuning of Association Cortex in Human Structural Brain Networks. Cereb Cortex. 28, 281–294 (2018).

33. S. Dehaene, J.-P. Changeux, Experimental and theoretical approaches to conscious processing. Neuron. 70, 200–227 (2011).

34. R. S. Desikan, F. Ségonne, B. Fischl, B. T. Quinn, B. C. Dickerson, D. Blacker, R. L. Buckner, A. M. Dale, R. P. Maguire, B. T. Hyman, M. S. Albert, R. J. Killiany, An automated labeling system for subdividing the human cerebral cortex on MRI scans into gyral based regions of interest. Neuroimage. 31, 968–980 (2006).

35. Y. Yeshurun, M. Nguyen, U. Hasson, Amplification of local changes along the timescale processing hierarchy. Proc Natl Acad Sci U S A. 114, 9475–9480 (2017).

36. R. E. Passingham, K. E. Stephan, R. Kötter, The anatomical basis of functional localization in the cortex. Nat Rev Neurosci. 3, 606–616 (2002).

37. H.-Y. S. Chien, C. J. Honey, Constructing and Forgetting Temporal Context in the Human Cerebral Cortex. Neuron. 106, 675-686.e11 (2020).

38. G. Shafiei, R. D. Markello, R. V. de Wael, B. C. Bernhardt, B. D. Fulcher, B. Mišić, Topographic gradients of intrinsic dynamics across neocortex. 178, 540–17 (2020).

39. A. Fornito, A. Zalesky, E. T. Bullmore, Network scaling effects in graph analytic studies of human resting-state FMRI data. Front Syst Neurosci. 4, 22 (2010).

40. A. Zalesky, A. Fornito, I. H. Harding, L. Cocchi, M. Yücel, C. Pantelis, E. T. Bullmore, Whole-brain anatomical networks: does the choice of nodes matter? Neuroimage. 50, 970–983 (2010).

41. Y. Liu, A. Beyer, R. Aebersold, On the Dependency of Cellular Protein Levels on mRNA Abundance. Cell. 165, 535–550 (2016).

42. R. D. Hodge, T. E. Bakken, J. A. Miller, K. A. Smith, E. R. Barkan, L. T. Graybuck, J. L. Close, B. Long, N. Johansen, O. Penn, Z. Yao, J. Eggermont, T. H. öllt, B. P. Levi, S. I. Shehata, B. Aevermann, A. Beller, D. Bertagnolli, K. Brouner, T. Casper, C. Cobbs, R. Dalley, N. Dee, S.-L. Ding, R. G. Ellenbogen, O. Fong, E. Garren, J. Goldy, R. P. Gwinn, D. Hirschstein, C. D. Keene, M. Keshk, A. L. Ko, K. Lathia, A. Mahfouz, Z. Maltzer, M. McGraw, T. N. Nguyen, J. Nyhus, J. G. Ojemann, A. Oldre, S. Parry, S. Reynolds, C. Rimorin, N. V. Shapovalova, S. Somasundaram, A. Szafer, E. R. Thomsen, M. Tieu, G. Quon, R. H. Scheuermann, R. Yuste, S. M. Sunkin, B. Lelieveldt, D. Feng, L. Ng, A. Bernard, M. Hawrylycz, J. W. Phillips, B. Tasic, H. Zeng, A. R. Jones, C. Koch, E. S. Lein, Conserved cell types with divergent features in human versus mouse cortex. Nature, 1–38 (2019).

43. S. Oldham, A. Arnatkevic Iūtė, R. E. Smith, J. Tiego, M. A. Bellgrove, A. Fornito, The efficacy of different preprocessing steps in reducing motion-related confounds in diffusion MRI connectomics. Neuroimage. 222, 117252 (2020).

44. J. L. R. Andersson, M. S. Graham, E. Zsoldos, S. N. Sotiropoulos, Incorporating outlier detection and replacement into a non-parametric framework for movement and distortion correction of diffusion MR images. NeuroImage. 141, 556–572 (2016).

45. J. L. R. Andersson, M. S. Graham, I. Drobnjak, H. Zhang, N. Filippini, M. Bastiani, Towards a comprehensive framework for movement and distortion correction of diffusion MR images: Within volume movement. NeuroImage. 152, 450–466 (2017).

46. Y. Zhang, M. Brady, S. M. Smith, Segmentation of brain MR images through a hidden Markov random field model and the expectation-maximization algorithm. IEEE Transactions on Medical Imaging. 20, 45–57 (2001).

47. J. D. Tournier, R. Smith, D. Raffelt, R. Tabbara, T. Dhollander, M. Pietsch, D. Christiaens, B. Jeurissen, C.-H. Yeh, A. Connelly, MRtrix3: A fast, flexible and open software framework for medical image processing and visualisation. NeuroImage, 116137 (2019).

48. J. D. Tournier, F. Calamante, A. Connelly, Robust determination of the fibre orientation distribution in diffusion MRI: Non-negativity constrained super-resolved spherical deconvolution. NeuroImage. 35, 1459–1472 (2007).

49. J.-D. Tournier, F. Calamante, D. G. Gadian, A. Connelly, Direct estimation of the fiber orientation density function from diffusion-weighted MRI data using spherical deconvolution. Neuroimage. 23, 1176–1185 (2004).

50. R. E. Smith, J.-D. Tournier, F. Calamante, A. Connelly, SIFT2: Enabling dense quantitative assessment of brain white matter connectivity using streamlines tractography. Neuroimage. 119, 338–351 (2015).

51. R. E. Smith, J. D. Tournier, F. Calamante, A. Connelly, Anatomically-constrained tractography: Improved diffusion MRI streamlines tractography through effective use of anatomical information. NeuroImage. 62, 1924–1938 (2012).

52. R. E. Smith, J. D. Tournier, F. Calamante, A. Connelly, The effects of SIFT on the reproducibility and biological accuracy of the structural connectome. NeuroImage. 104, 253–265 (2015).

53. J. A. Roberts, A. Perry, G. Roberts, P. B. Mitchell, M. Breakspear, Consistency-based thresholding of the human connectome. Neuroimage. 145, 118–129 (2017).

54. K. Sabaroedin, J. Tiego, L. Parkes, F. Sforazzini, A. Finlay, B. Johnson, A. Pinar, V. Cropley, B. J. Harrison, A. Zalesky, C. Pantelis, M. Bellgrove, A. Fornito, Functional Connectivity of Corticostriatal Circuitry and Psychosis-like Experiences in the General Community. Biol Psychiatry. 86, 16–24 (2019).

55. S. M. Smith, M. Jenkinson, M. W. Woolrich, C. F. Beckmann, T. E. J. Behrens, H. Johansen-Berg, P. R. Bannister, M. De Luca, I. Drobnjak, D. E. Flitney, R. K. Niazy, J. Saunders, J. Vickers, Y. Zhang, N. De Stefano, J. M. Brady, P. M. Matthews, Advances in functional and structural MR image analysis and implementation as FSL. Neuroimage. 23, S208–S219 (2004).

56. L. Griffanti, G. Salimi-Khorshidi, C. F. Beckmann, E. J. Auerbach, G. Douaud, C. E. Sexton, E. Zsoldos, K. P. Ebmeier, N. Filippini, C. E. Mackay, S. Moeller, J. Xu, E. Yacoub, G. Baselli, K. Ugurbil, K. L. Miller, S. M. Smith, ICA-based artefact removal and accelerated fMRI acquisition for improved resting state network imaging. Neuroimage. 95, 232–247 (2014).

57. B. B. Avants, N. J. Tustison, G. Song, P. A. Cook, A. Klein, J. C. Gee, A reproducible evaluation of ANTs similarity metric performance in brain image registration. Neuroimage. 54, 2033–2044 (2011).

58. K.-F. Wong, X.-J. Wang, A recurrent network mechanism of time integration in perceptual decisions. J Neurosci. 26, 1314–1328 (2006).

59. L. F. Abbott, F. S. Chance, Drivers and modulators from push-pull and balanced synaptic input. Prog Brain Res. 149, 147–155 (2005).

60. K. E. Stephan, N. Weiskopf, P. M. Drysdale, P. A. Robinson, K. J. Friston, Comparing hemodynamic models with DCM. Neuroimage. 38, 387–401 (2007).

61. V. S. Sohal, J. L. R. Rubenstein, Excitation-inhibition balance as a framework for investigating mechanisms in neuropsychiatric disorders. Mol Psychiatry. 24, 1248–1257 (2019).

62. J. H. Krystal, A. Anticevic, G. J. Yang, G. Dragoi, N. R. Driesen, X.-J. Wang, J. D. Murray, Impaired Tuning of Neural Ensembles and the Pathophysiology of Schizophrenia: A Translational and Computational Neuroscience Perspective. Biol Psychiatry. 81, 874–885 (2017).

63. J. Arloth, D. M. Bader, S. Röh, A. Altmann, Re-Annotator: Annotation Pipeline for Microarray Probe Sequences. PLoS One. 10, e0139516 (2015).

64. J. A. Miller, V. Menon, J. Goldy, A. Kaykas, C.-K. Lee, K. A. Smith, E. H. Shen, J. W. Phillips, E. S. Lein, M. J. Hawrylycz, Improving reliability and absolute quantification of human brain microarray data by filtering and scaling probes using RNA-Seq. BMC Genomics. 15, 154 (2014).

65. J. M. Henley, K. A. Wilkinson, Synaptic AMPA receptor composition in development, plasticity and disease. Nat Rev Neurosci. 17, 337–350 (2016).

66. P. Paoletti, C. Bellone, Q. Zhou, NMDA receptor subunit diversity: impact on receptor properties, synaptic plasticity and disease. Nat Rev Neurosci. 14, 383–400 (2013).

67. E. Sigel, M. E. Steinmann, Structure, function, and modulation of GABA(A) receptors. J Biol Chem. 287, 40224–40231 (2012).

68. S. Vicini, J. F. Wang, J. H. Li, W. J. Zhu, Y. H. Wang, J. H. Luo, B. B. Wolfe, D. R. Grayson, Functional and pharmacological differences between recombinant N-methyl-D-aspartate receptors. J Neurophysiol. 79, 555–566 (1998).

69. G. Deco, J. Cruzat, J. Cabral, G. M. Knudsen, R. L. Carhart-Harris, P. C. Whybrow, N. K. Logothetis, M. L. Kringelbach, Whole-Brain Multimodal Neuroimaging Model Using Serotonin Receptor Maps Explains Non-linear Functional Effects of LSD. Curr Biol. 28, 3065-3074.e6 (2018).

70. M. L. Kringelbach, J. Cruzat, J. Cabral, G. M. Knudsen, R. Carhart-Harris, P. C. Whybrow, N. K. Logothetis, G. Deco, Dynamic coupling of whole-brain neuronal and neurotransmitter systems. Proceedings of the National Academy of Sciences. 117, 9566– 9576 (2020).

71. M. F. Glasser, D. C. Van Essen, Mapping human cortical areas in vivo based on myelin content as revealed by t1-and t2-weighted MRI. Journal of Neuroscience. 31, 11597– 11616 (2011).

72. F. Váša, R. Romero-Garcia, M. G. Kitzbichler, J. Seidlitz, K. J. Whitaker, M. M. Vaghi, P. Kundu, A. X. Patel, P. Fonagy, R. J. Dolan, P. B. Jones, I. M. Goodyer, the NSPN Consortium, P. E. Vértes, E. T. Bullmore, Conservative and disruptive modes of adolescent change in brain functional connectivity. bioRxiv, 1–8 (2019).

73. A. Fornito, Fundamentals of Brain Network Analysis (Academic Press, Inc, London, 2016).

74. F. Váša, R. Romero-Garcia, M. G. Kitzbichler, J. Seidlitz, K. J. Whitaker, M. M. Vaghi, P. Kundu, A. X. Patel, P. Fonagy, R. J. Dolan, P. B. Jones, I. M. Goodyer, NSPN Consortium, P. E. Vértes, E. T. Bullmore, Conservative and disruptive modes of adolescent change in human brain functional connectivity. Proceedings of the National Academy of Sciences. 117, 3248–3253 (2020).

